# Second-order effects of chemotherapy pharmacodynamics and pharmacokinetics on tumor regression and cachexia

**DOI:** 10.1101/2023.06.14.544974

**Authors:** Luke Pierik, Patricia McDonald, Alexander R. A. Anderson, Jeffrey West

**Affiliations:** Center for Complex Biological Systems, University of California Irvine, Irvine, CA; Department of Cancer Physiology, Moffitt Cancer Center, Tampa, FL; Integrated Mathematical Oncology, Moffitt Cancer Center, Tampa, FL

## Abstract

Drug dose response curves are ubiquitous in cancer biology, but these curves are often used to measure differential response in first-order effects: the effectiveness of increasing the cumulative dose delivered. In contrast, second-order effects (the variance of drug dose) are often ignored. Knowledge of second-order effects may improve the design of chemotherapy scheduling protocols, leading to improvements in tumor response without changing the total dose delivered. By considering treatment schedules with identical cumulative dose delivered, we optimize treatment by comparing high variance schedules (e.g. high dose, low dose) with low variance schedules (constant dose). We extend a previous framework used to quantify second-order effects, known as antifragility theory, to investigate the role of drug pharmacokinetics. Using a simple one-compartment model, we find that high variance schedules are effective for a wide range of cumulative dose values. Next, using a mouse-parameterized two-compartment model of 5-fluorouracil, we show that the optimal schedule depends on initial tumor volume. Finally, we illustrate the trade-off between tumor response and lean mass preservation. Mathematical modeling indicates that high variance dose schedules provide a potential path forward in mitigating the risk of chemotherapy-associated cachexia by preserving lean mass without sacrificing tumor response.

## 1 Introduction

In probability theory, the first moment of a probability distribution is known as the mean of the underlying distribution and the second moment is the variance. Thus, so-called “first-order effects” relate the response of a system to altering the input mean while “second-order effects” relate the system’s response to the input variance. A mathematical framework used to quantify these so-called second-order effects is known as antifragile theory^1^. The term antifragile was coined to describe systems that benefit from volatility in the input distribution, serving as a contrast to fragile systems that are harmed by input volatility. Thus far, the concept has been heavily applied to financial markets, where an individual’s investment portfolio may be susceptible to market volatility (fragile), leading them to financial ruin^2^. In contrast, it’s possible to design an investment strategy based on the premise of deriving financial *benefit* from volatile fluctuations in market prices.

These concepts, rooted in probability theory, can be readily translated into oncology^3^. For example, cancer chemotherapy is intended to induce dramatic perturbations to a tumor, leading to increased cell death. The timing of these perturbations is determined by a treatment scheduling protocol. By definition, each chemotherapy treatment scheduling protocol has an associated mean and variance. Thus, we hypothesize that treatment design should account for both first-order (optimizing the mean of the dose delivered) and second-order (optimizing the variance of the dose delivered) effects. Historically, the design of chemotherapeutic schedules was intended to maximize the first-order effect, typically through a “maximum-tolerable dose” (MTD) paradigm. Improving a first-order effect is achieved either through increasing the cumulative dose delivered^4^ or shortening the time between subsequent doses^5^. In order to improve the design of chemotherapy scheduling, we propose a strategy to quantify and optimize second-order effects, which consider the dose variance, in addition to the dose mean.

### 1.1 Antifragility in cancer

Mathematical modeling is highly suitable to quantify second-order effects and to predict the effectiveness of drug scheduling protocols^3,6,7^. Previously, we introduced a framework for “antifragile therapy” wherein the curvature of the dose response curve determines the optimal treatment schedule^8^. In this initial work, we considered two treatment protocols: 1) even dosing: continual administration of the same dose at regular intervals, and 2) uneven dosing: a high dose followed by a low dose, in alternating fashion. Examples of this bolus dosing are shown in figure 1A. Assuming no drug decay (figure 1B), the optimal schedule can be determined by observing the shape (curvature) of the dose response function (figure 1C).

**Figure 1.**
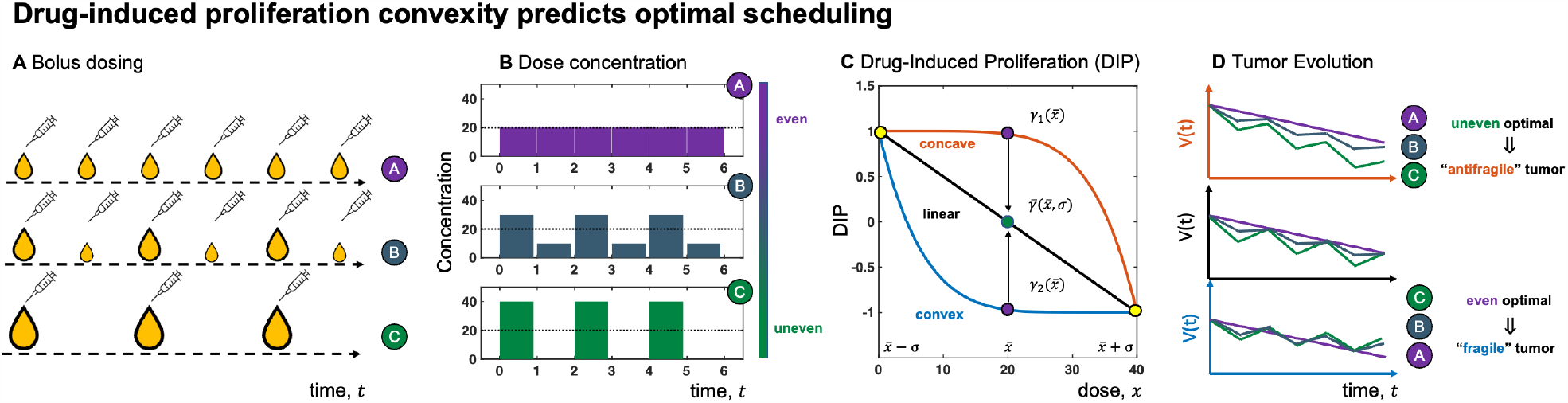
Drug-induced proliferation convexity predicts optimal scheduling. (A,B) Schematics of different dose schedules from even to uneven treatment with same cumulative dose delivered. (C) Examples of drug-induced proliferation (DIP) curves with differing convexity. Green dot indicates difference between the average DIP for two doses versus the DIP at the average dose delivered. (D) Predicted tumor evolution based on the convexity of the DIP curve. When the DIP curve is convex, more tumor is eliminated and the tumor is antifragile. Inversely, if the DIP curve is concave, even treatment is optimal and the tumor is antifragile.

In the event that a dose response curve is convex, even dosing is optimal (figure 1D). Conversely, if the underlying dose response is concave, uneven dosing is optimal: a high dose followed by a low dose. Others have followed similar approaches in applying antifragile theory to cancer applications including immune-tumor interactions^9^, biological boolean networks^10^, molecular biology and evolution^11^.

A major limitation of previous work applying antifragility to treatment scheduling has been the lack of pharmacokinetic considerations. Our previous work^3,8,12^ has considered drug pharmacodynamics in isolation, where the curvature of dose response curves may provide predictive value in a limited preclinical (e.g. in vitro) or theoretical context. Figure 1 illustrates the predictive power of this graphical approach that links dose response convexity and concavity to outcomes, but this graphical approach is confounded by the drug’s pharmacokinetic decay and absorption before the drug reaches the target tumor. To rectify this limitation, we introduce a mathematical model of tumor growth, response, and chemotherapy-induced cachexia as a test-bed for quantifying the second-order effects of treatment. Here, we go beyond previous results to include both drug pharmacodynamics (PD) and pharmacokinetics (PK) to quantify the second-order effects of chemotherapy on tumor regression.

To do so, we begin with a simple, single-compartment PK/PD model to illustrate the influence of PK decay rates on emergent second-order treatment effects. We introduce a metric that describes tumor fragility (or antifragility) that predicts whether even or uneven dosing achieves optimal tumor regression. In the subsequent sections, we apply this metric to a previously published mouse-parameterized two-compartment PK/PD model of 5-fluorouracil^13,14^. This model allows us to explore the effect of an additional pharmacokinetic compartment on tumor fragility. Importantly, the model also considers a potential side effect of chemotherapy through cachexia.

### 1.2 Chemotherapy-induced cachexia

Cancer-associated cachexia, defined as the progressive wasting and weakness of skeletal muscle with or without fat loss^15^ is a debilitating complication of cancer that reduces quality of life and response to anticancer therapies compromising survival. However, both preclinical^16,17^ and clinical studies^18–20^ have shown that anticancer therapies themselves can promote or exacerbate cachexia. Often, there is a trade-off between tumor regression and chemotherapy-induced side effects, where dose escalation may result in increased tumor regression, but at the risk of increased patient toxicity or other side effects including skeletal muscle atrophy and weakness; a specific off-target systemic effect that manifests clinically as cachexia. Hence chemo-induced cachexia has recently emerged as a critical clinical problem.

In the second half of this manuscript, we quantify the second-order treatment effects on both tumor regression and lean mass. Second-order effects have not, to our knowledge, been quantified in cachexia and remains an unexplored potential avenue for reducing the rates of cachexia in patients receiving chemotherapy. For example, the first-order effect of increasing mean dose has an adverse effect on increasing the likelihood of cachexia. In this manuscript, we advocate for the quantification of second-order effects so that both the mean dose and the dose variance can be optimized, in favor of reducing tumor burden and maintaining lean mass to avoid cachexia. Optimizing second-order effects may provide a potential path forward in mitigating the risk of cancer cachexia by preserving lean mass without sacrificing tumor response.

### 1.3 The mathematics of antifragility

We consider a suite of dosing schedules like those in panel A in figure 1, where a pair of doses is administered over a time interval *T* . This is repeated for *n* cycles (here *n* = 6), where each cycle has the same cumulative dose. Therefore, the total drug delivered to the patient is the same for schedules A, B, and C. Treatment schedules A, B, and C are characterized by a dosing vector 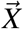 with 2-dose cycles governed by a prescribed dose variance *σ* and denoted as 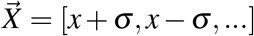 for *n* cycles. As shown, schedules A, B, and C transition from purple to green indicating an increase in *σ*.

The drug-induced proliferation (DIP, or symbolically *γ*) rate determines the pharmacodynamics of tumor killing. DIP represents the steady state proliferation rate of cells in the presence of a drug^21–23^, and thus tumors are decaying if DIP is less than zero. Optimal treatment would minimize DIP, hence maximizing tumor regression. As shown in figure 1C, its shape can determine whether uneven or even treatment gives a higher effective *γ*. In particular, the concave *γ* in figure 1C predicts an uneven schedule is more beneficial (the green dot representing the average of high and low dose is below the purple dot representing the instantaneous dose). The convex *γ* indicates even treatment would be optimal. Finally, a linear DIP curve leads to identical outcomes for even and uneven treatment (no advantage).

Panel D illustrates examples for all three scenarios under different treatment schedules described earlier. Under a convex DIP, we have uneven treatment being optimal for tumor suppression, so we say the system is antifragile (benefiting from dose volatility) while the concave DIP predicts even treatment is more beneficial, so the system is fragile (benefiting from dose consistency). This motivates the definition of fragility in cancer:

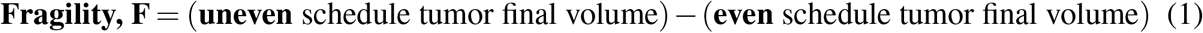

Fragility provides a quantifiable metric delineating when an uneven schedule or even schedule is optimal. If fragility is negative (*F <* 0) an uneven schedule minimizes the final tumor volume and is optimal. We term this situation antifragile (figure 1D, top). If fragility is positive (*F >* 0) an even schedule is optimal. We term this situation fragile (figure 1D, bottom). If fragility is zero, then neither schedule outperforms the other (figure 1D, middle). In the subsequent sections below, we focus on defining the boundary between fragile and antifragile regions in parameter space.

## 2 Methods (Model development)

### 2.1 Fragility for daily dosing

As stated above, we employ antifragile theory to compare the outcomes between even and uneven dosing schedules. A treatment cycle is defined as two sequential doses, 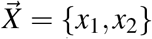 Here, we restrict ourselves to treatment cycles with identical mean dose, 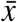, such that 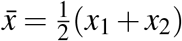. We can define all possible such schedules with two control parameters, 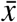 and *σ*:

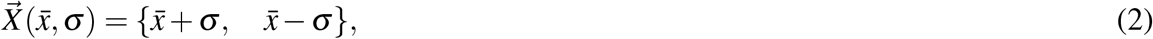

where 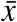 is the mean dose and *σ* is the dose variance. Note that *σ* = 0 corresponds to an even treatment schedule.

To evaluate second-order effects, we consider the following “naive” definition of fragility, motivated by Jensen’s inequality^24^. See supplemental section S1 for more discussion on Jensen’s inequality. We assume that the effect of each dose is independent and thus fragility is the difference between the average volume resulting from a high and low dose, 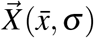, and the average of two even doses, 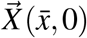:

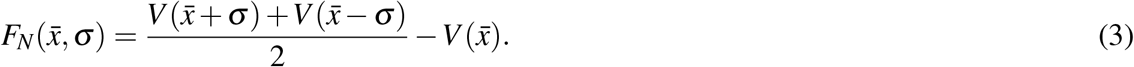

An uneven schedule is more effective against the tumor as the even schedule when:

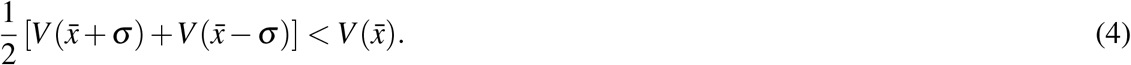

This expression implies that the function *V* (*y*) is concave in the interval 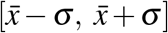 according to Jensen’s Inequality. Conversely, the following expression implies the function *V* (*y*) is convex:

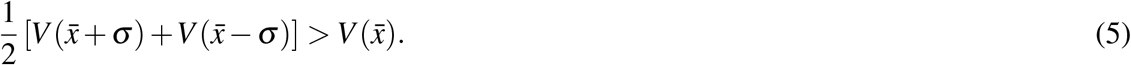

This is equivalent to stating *F <* 0 (antifragile; concave) or *F >* 0 (fragile; convex). We refer to equation 3 as the naive form of fragility, denoted by the subscript *N, F*_*N*_, because it implicitly assumes path-independence (the ordering of high and low dose does not matter). In subsequent sections we will show that this assumption cannot be made. Thus, for the rest of the manuscript we will use the following definition for fragility, based on the difference in final outcome of each treatment schedule:

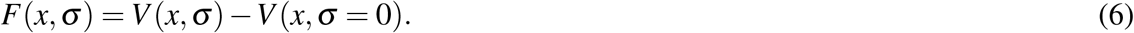

### 2.2 Fragility for weekly dosing

Next, we generalize the range of possible treatment schedules to consider a fixed number of *N* doses (e.g. daily administration for a week, *N* = 7). Within these *N* doses, we administer a fixed number of doses, *x*, followed by *H* treatment holidays, *x* = 0

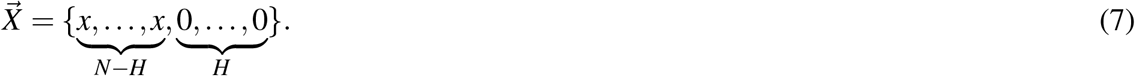

In this way, we compare weekly (*N* = 7) dosing schedules that range from even administration (*H* = 0) with a daily dose of 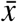, compared to uneven administration of *N − H* doses, *x*′, followed by *H* number of zero dose holidays, *x* = 0. Again, we restrict the set of dosing schedules to identical cumulative dose 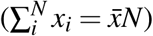, allowing us to solve for the high dose, *x*′, as a function of the number of holidays *H*:

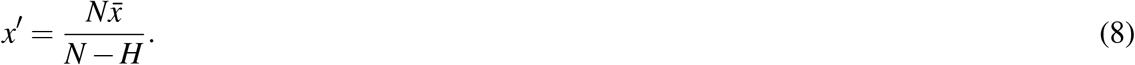

This restriction encompasses commonly tested chemotherapy regimens. For example, below we re-purpose data from Farhang-Sardroodi et. al.^14^ (see Results) that compares daily administration of 5-fluorouracil (*H* = 0) to a (5,2) schedule of five daily doses followed by two holidays (*H* = 2). We thus compare final tumor volumes between an even treatment and uneven treatment through the following equation for fragility:

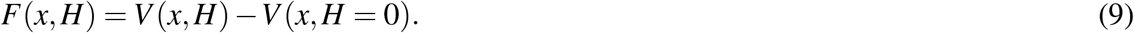

### 2.3 1-Compartment Model

Next, we apply this definition of fragility to a 1-compartment model pharmacokinetic model of drug concentration, *x*(*t*), coupled to our DIP pharmacodynamic model of tumor growth and decay, *γ*(*x*). The tumor is exponentially growing (or decaying) according to DIP rate, *γ*, as a function of the dose, *x*. Let *V* (*t*) be a tumor volume at time *t* (treatment ends at *t* = *T*) and *γ*(*x*) be the DIP growth rate of the tumor. We then have the following expression for tumor volume:

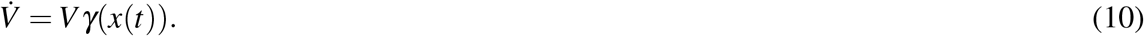

where *x*(*t*) is the instantaneous drug concentration near the tumor at time *t*. In practice, *γ*(*x*) can be expressed as the Hill function, a commly used empirical pharmacodynamic equation^21,25,26^:

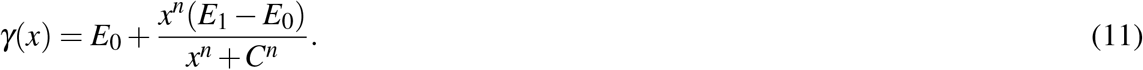

Given the treatment dynamics, *x*(*t*), and initial conditions, we can solve for the tumor volume over time. This tumor volume can be used to evaluate the fragility of our tumor in response to different treatment schedules. When analyzing the fragility of the system, we can explicitly consider the influence of pharmacokinetics; if *x*(*t*) does not decay, then the schedule is piecewise constant over the dosing interval [0, *T*] and pharmacokinetic effects are ignored (e.g. figure 1B).

This Hill function typically has both convex and concave regions of the dose response curve, separated by inflection point. The inflection point can be found analytically by finding the dose *x*^*^ where the second-derivative is zero (e.g. see ref.^8^), and it is given by:

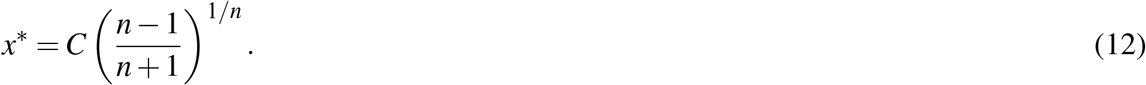

### 2.4 2-compartment PK model of tumor response and cachexia

It is common practice to extend pharmacokinetics to consider two compartments: drug concentration in the plasma, *x*_1_(*t*), and in the tissue, *x*_2_(*t*). In this manuscript, we explore a previously published two-compartment PK/PD model by Farhang-Sardroodi et al parameterized by an in vivo study to determine the utility of fragility^14^. Of note is the two-compartment model for drug transfer between plasma (*x*_1_) and tissue (*x*_2_), representing model 7 of Win-NonLin^27^. The resulting tumor growth law is given by an underlying exponential-linear growth with decreasing factor based on tissue concentration:

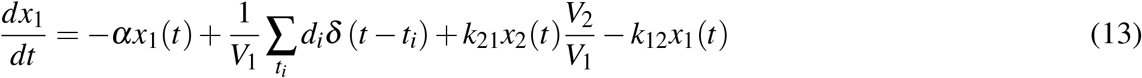

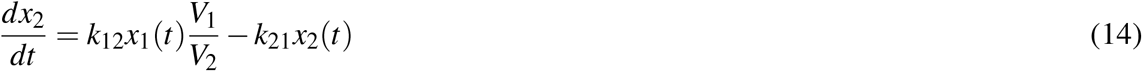

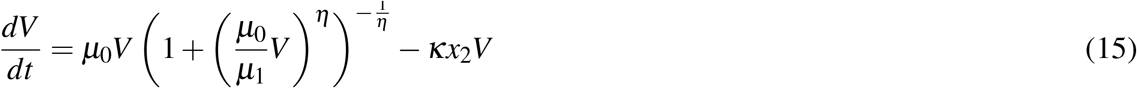

Instantaneous injections of a drug are given by the sum of Dirac delta, *δ* (*t − t*_*i*_), at treatment administration time points *t*_*i*_. Here, *α* continues to act as the drug clearance variable while *k*_12_ and *k*_21_ represent the transition rates between the plasma and tissue compartments, respectively. Finally, *κ* is a rate of tumor death per unit tissue-muscle volume, scaled by the drug concentration in the tissue, *x*_2_(*t*).

## 3 Results

### 3.1 1-Compartment Model, without pharmacokinetics

Using the 1-compartment model described earlier, we begin with the simple case of a single cycle, a pair of doses administered in the time interval *T* . The time-dependent dose concentration *x*(*t*) is given by:

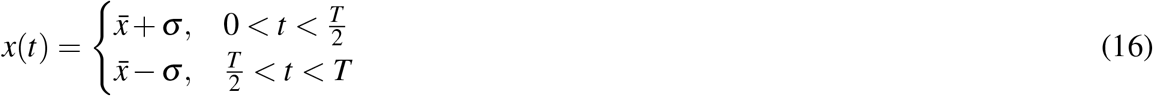

where *σ* is a parameter determining the unevenness of the dosing schedule and 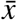 is the mean dose experienced by the tumor 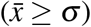. Note that in this case there is no decay in drug concentration between dosing events, so we are in a case without pharmacokinetics.

Inserting *x*(*t*) into the tumor volume equation 38, we can derive an expression for *V* (*t*) which only depends on our DIP curve, *γ*:

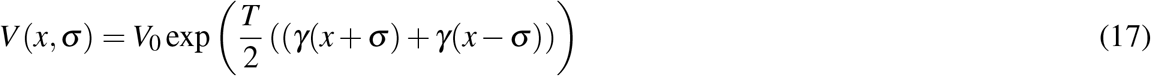

Given this schedule, we can describe a corresponding fragility defined as the difference in final tumor size for uneven and even treatment after an interval *T* :

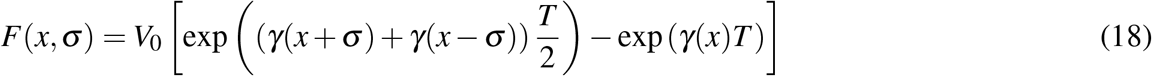

The tumor dynamics and fragility resulting from this setup are presented in Panels A-E in Figure 2. Panel A shows the model’s compartments with the treatment dose directly influencing tumor volume. Panel B illustrates the even (*σ* = 0) schedule in purple and uneven 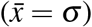 schedule in green, both without PK clearance. The schedules in Panel B create tumor trajectories according to Panel C, where the even treatment produces a smaller tumor than the uneven treatment. Panel D shows an example hypothetical DIP curve determining tumor growth that has a characteristic sigmoidal pattern parameterized from the Hill function. The interaction between the DIP curve and treatment schedule is shown in Panel E, where the fragility is given over a range of average doses 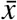 and schedules (see supplemental section S2 for how this compares to previous work^8^). Here, the concave region of the DIP curve in D corresponds to an antifragile tumor and the convex region of the DIP curve corresponds to a fragile tumor. Thus, uneven is optimal for low doses (*F <* 0) while even is optimal for high doses (*F >* 0).

**Figure 2.**
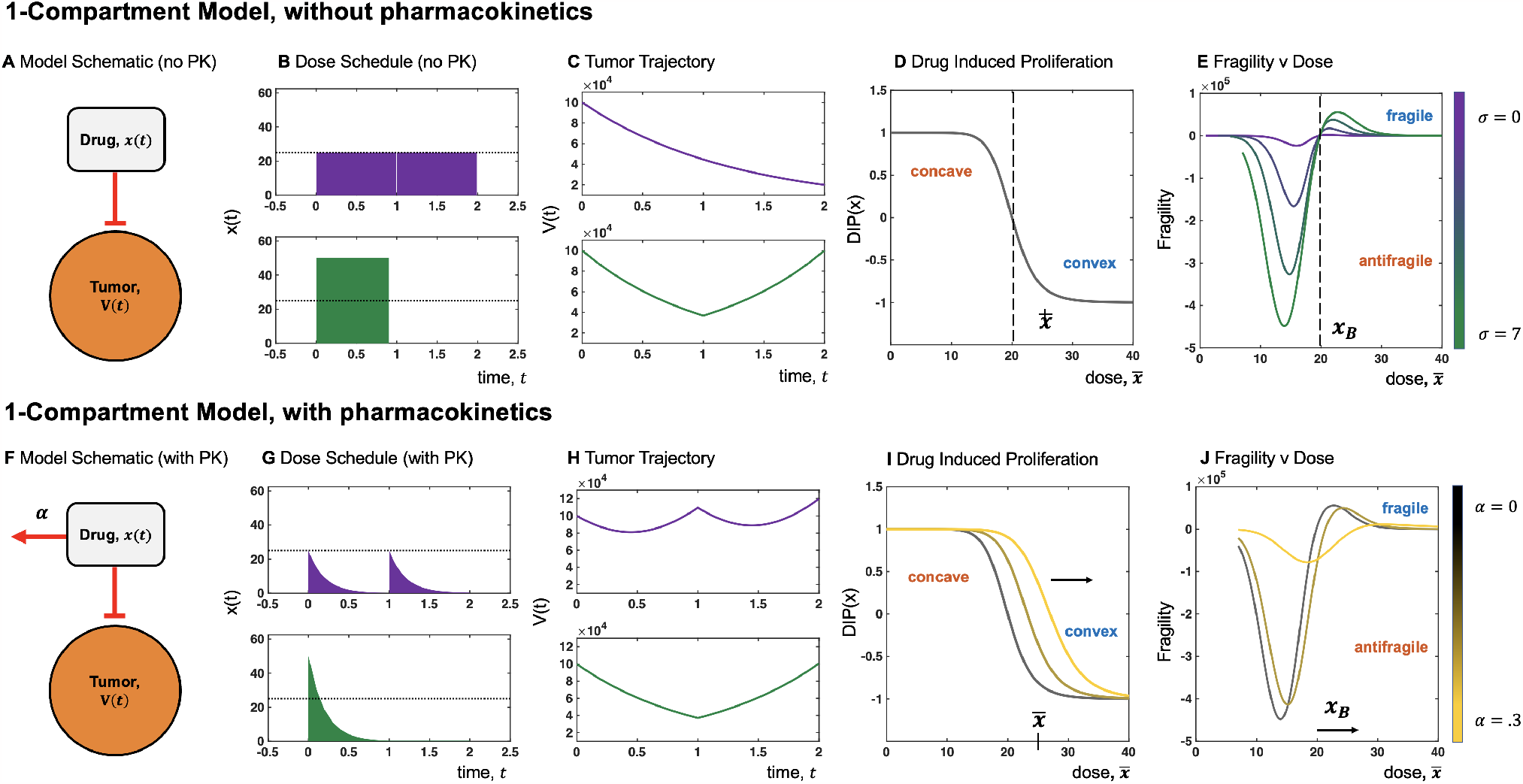
Pharmacokinetic-induced fragility in a 1-Compartment model. (A,F) 1-compartment model setup for drug delivery and tumor elimination with and without pharmacokinetic decay. (B,G) Schematic dose schedules for even and uneven treatments with and without pharmacokinetics for 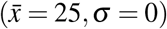 and 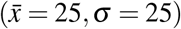 treatment cycles. (C,H) Example tumor trajectories resulting from the dosing schedules in (B,G); while tumor size is smaller following even treatment in (C), there is a smaller tumor size following uneven treatment in (H). (D,I) A DIP curve for a tumor and its dependence on treatment decay. For *α* = [0 .15 .3], increasing pharmacokinetics correspond to an increased region of concavity. (E) Fragility values across a range of average doses, *x* and variance *σ* = [1 3 5 7]; (J) Fragility values across a range of average doses *x* and pharmacokinetics *α* = [0 .15 .3]. Move from antifragile to fragile treatment as *x* increases in both (E) and (J), but as *α* increases, the range of doses which give an antifragile tumor increases. For all figures, the Hill function parameters were *E*_0_ = 1, *E*_1_ = -1, *C* = 20, *n* = 10.

### 3.2 1-Compartment Model, with pharmacokinetic drug decay

To consider the role of pharmacokinetics in the 1-compartment model, we now consider a similar single-cycle schedule where the initial drug concentration undergoes exponential decay. This model follows the structure of model 1 from Win-NonLin^27^ with one-compartment, bolus input, and first-order output (figure 2F). If we assume that there is negligible residual concentration of drug prior to the second half of the cycle, we have the following expression for dose concentration:

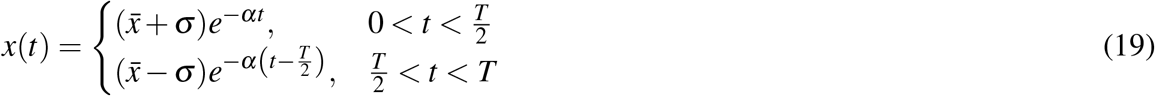

Panels F-J in Figure 2 illustrate the one-compartment model with decay and how it influences treatment outcomes. Panel F and Panel G show the introduction of decay and the resulting exponential decay in drug dose experienced by the tumor. Panel H shows the resulting tumor trajectories, with the uneven schedule optimal eliminating more tumor than the even schedule. Panel I shows that as the *α* decay parameter increases, the DIP curve shifts to the right, increasing the region of concavity. This corresponds to the shifting fragility in Panel J, where increasing PK clearance rate, *α*, corresponds to an extended antifragile treatment regime (black to yellow lines).

Introducing pharmacokinetics to this 1-compartment model alters the implications of treatment in two significant ways. First, the antifragile-fragile boundary increases as the decay parameter *α* increases (*x*_*B*_ in figure 2J). The new relationship between the decay and threshold concentration, *x*_*B*_(*α*), is given as an implicit function demonstrated in figure 3, and there is a greater region where antifragile treatment scheduling is beneficial. This relationship can be explained by the fact that when drugs decay, they decay to a lower concentration, so in effect, the dynamics shown in the above figure may be seen as a shifting the constant-drug treatment shown earlier toward a higher “effectively low dose” threshold. Second, the intensity or degree of benefit and harm of using either fragile or antifragile treatment decreases as the decay rate increases. The magnitude of maximum (anti-)fragility in figure 2J is reduced for high values of *α*. This intuitively follows by acknowledging that if the decay rate is high, then the most beneficial aspects of the treatment from higher dosages will decrease more rapidly and thus destroy less cancer, diminishing the influence of a higher drug concentration.

**Figure 3.**
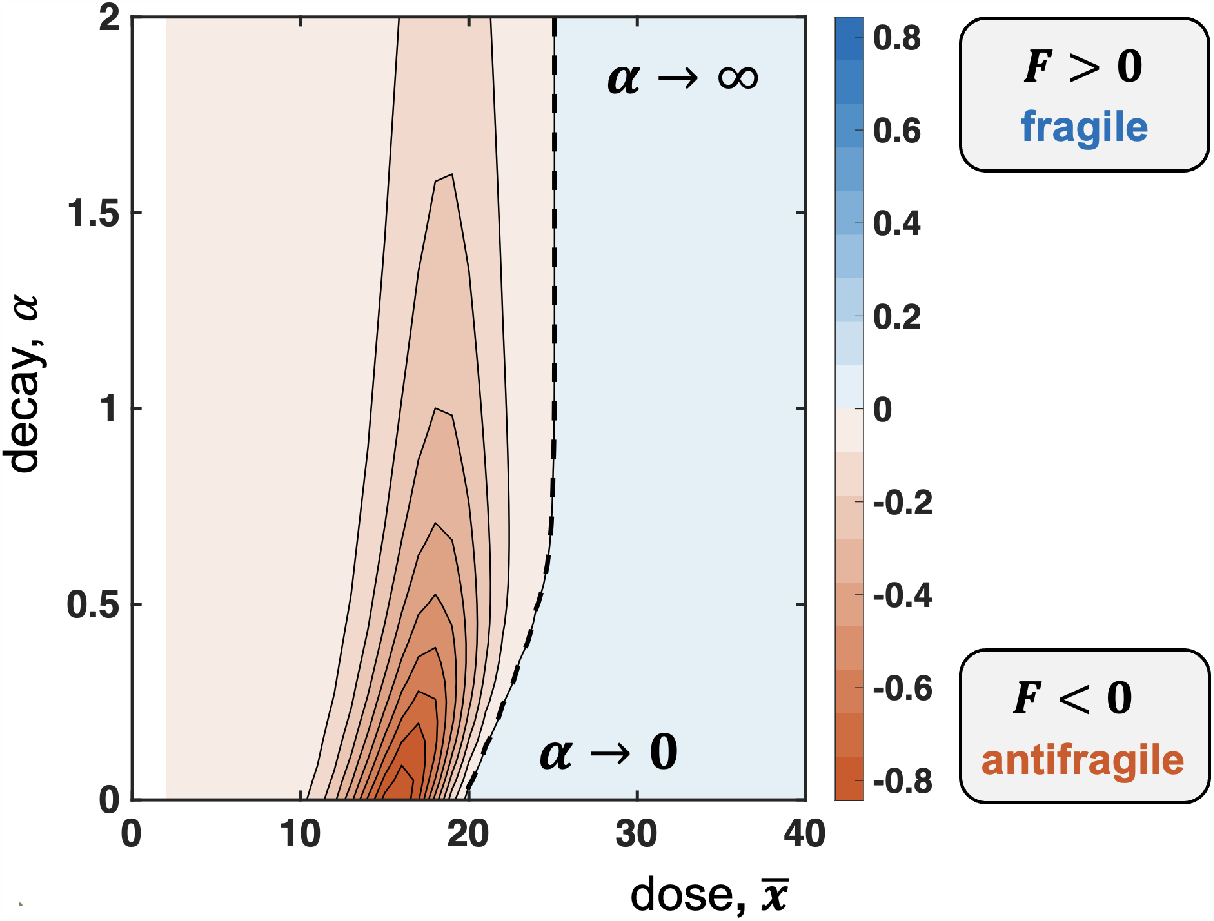
Pharmacokinetics and dose effect on fragility. When given an array of treatment doses and rates of drug decay, higher doses tend to be fragile while lower doses tend to be antifragile. The threshold between these optimal schedules changes as a function of *α*, with a lower bound determined by no pharmacokinetics (*α* → 0) and an upper bound determined by fast pharmacokinetics (*α* → ∞). Figure generated from equations 49 - 51 with *σ* = 2, *E*_0_ = 1, *E*_1_ = -1, *C* = 20, *n* = 10, and normalized by initial tumor volume.

#### 3.2.1 Variable pharmacokinetic decay (*α* 0)

Because the form of the DIP curve, *γ*(*x*), is explicitly defined (eqn. 11), an explicit form for the tumor volume can be obtained:

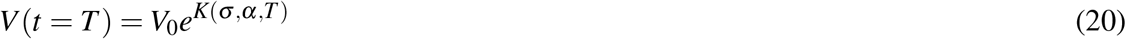

where

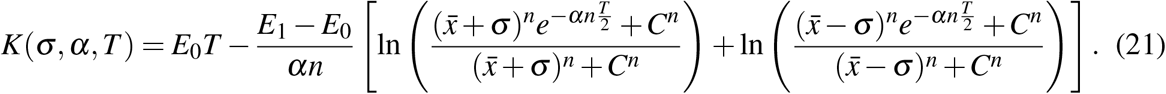

Because we know the tumor volume following a treatment cycle, we can obtain the corresponding fragility:

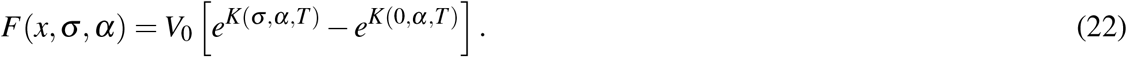

Equation 22 is an implicit function of the baseline dose 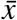 and drug decay rate, *α*, shown in figure 3 for a range of decay rate and dose values. We note two important cases for the implicit function: infinitely fast decay (*α* → ∞) and no drug decay (*α* = 0).

#### 3.2.2 Infinite pharmacokinetic decay (*α* → ∞)

In the case of fast pharmacokinetics, we also can retrieve an implicit function for the antifragile-fragile boundary by taking the limit (see section S3.2 for full derivation); it is given by:

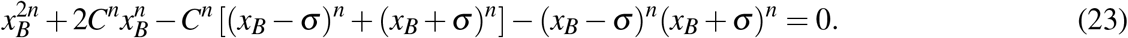

This *x*_*B*_ can be seen in figure 3, as the antifragile-fragile boundary (dashed line) is a straight vertical line for large values of *α*.

#### 3.2.3 No pharmacokinetic decay (*α* = 0)

When *α* = 0, equation 22 simply reduces to our earlier non-PK result in equation 17, so the tumor volume will be determined by:

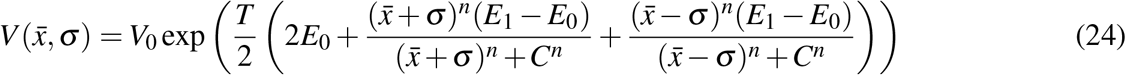

Where *γ*(*x*) is replaced by a parameterized Hill function (see section S3.3 for full derivation). In this case, the antifragile-fragile boundary can be analytically found when *σ*≈ 0 as

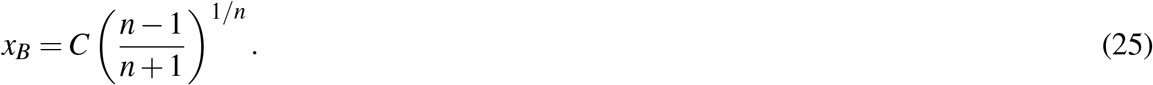

This result intuitively corresponds to the inflection point of the Hill function (eqn. 12), which delineates the concave region (low doses) from the convex (high doses) region of the dose response curve. More explanation can be found in refs. 3,8. Without pharmacokinetics, the antifragile-fragile boundary is only dependent on parameters from the DIP curve. This threshold corresponds to the far-left limiting value in figure 3. If we consider pharmacokinetic decay, this *x*_*B*_ also acts as a lower bound for the antifragile-fragile boundary.

### 3.3 The effect of residual dose

Thus far, we have made the assumption that the residual dose concentration is negligible at the end of each half cycle (e.g. 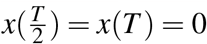). Revisiting this assumption, the dose concentration for a single treatment cycle is given by:

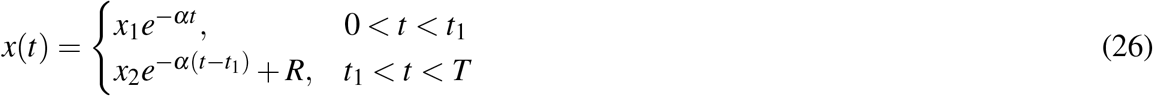

where the residual dose, *R*, is defined as:

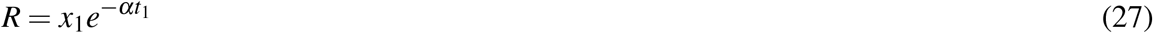

Note that if *R* ≈0, then we return to equation 19 for describing dose concentration dynamics. Intuitively, the residual dose concentration can be ignored in the case that PK clearance is very rapid (high *α*), the time between doses delivered is large (high *t*_1_), or the initial dose is low (small *x*_1_).

#### 3.3.1 High or low dose first

Thus far, we have only considered a high dose followed by a low dose: 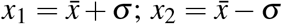. We now relax that assumption to consider the optimal dose ordering when residual dose is non-negligible. We find that when residual concentrations are non-negligible, results depend on the order in which high and low doses are delivered. The average amount of drug affecting the tumor will differ (see supplemental section S3.4). This results in a difference of first-order effects, given by:

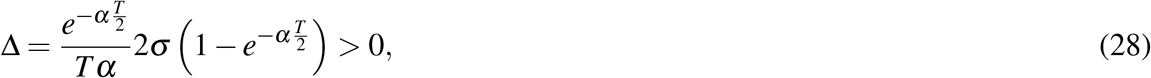

where Δ is the difference in total concentration delivered between high dose first and low dose first. It can be shown that Δ is a positive function (for *α, T, σ>* 0) and thus the high dose first always maximizes the total amount of drug delivered within the time span, *T* . Note: this difference decreases as a function of *α* and *T*, so the negligible concentration assumption is valid for fast pharmacokinetics or long time intervals between doses. See supplemental sections S3.1 and S3.4 for discussion on multiple treatment cycles and derivation of Δ.

### 3.4 2-Compartment PK Model Results

Next, we extend our investigation of the influence of pharmacokinetics on second-order effects by examining a two-compartment model of dose concentration in plasma and tissue. A schematic of the model is shown in figure 4A where the drug is intravenously delivered to a plasma compartment. The model accounts for plasma clearance at a rate *α* and transfer rates, *r*_*ij*_, between plasma and tissue compartments. The model tracks both the plasma concentration, *x*_1_(*t*), and tissue concentration, *x*_2_(*t*), over time. Tissue concentration has a direct effect on tumor kill (see eqns. 13 - 15 in Methods). Additionally, this system includes a lean mass compartment inhibited by the drug concentration in the tissue compartment. Therefore fragility can be quantified for both tumor response and lean mass. Panel B demonstrates the exponential-linear growth rate assumed for the tumor without treatment (see Methods). This mathematical model of chemotherapy (5-fluorouracil) was previously parameterized to mouse experiments^13,14^ by deriving a set of parameters to fit tumor volume data and lean mass data over time. We use this model and its associated parameterization as a point of departure for our purpose here to define the effect of pharmacokinetics on second-order effects in a biologically realistic mathematical model.

**Figure 4.**
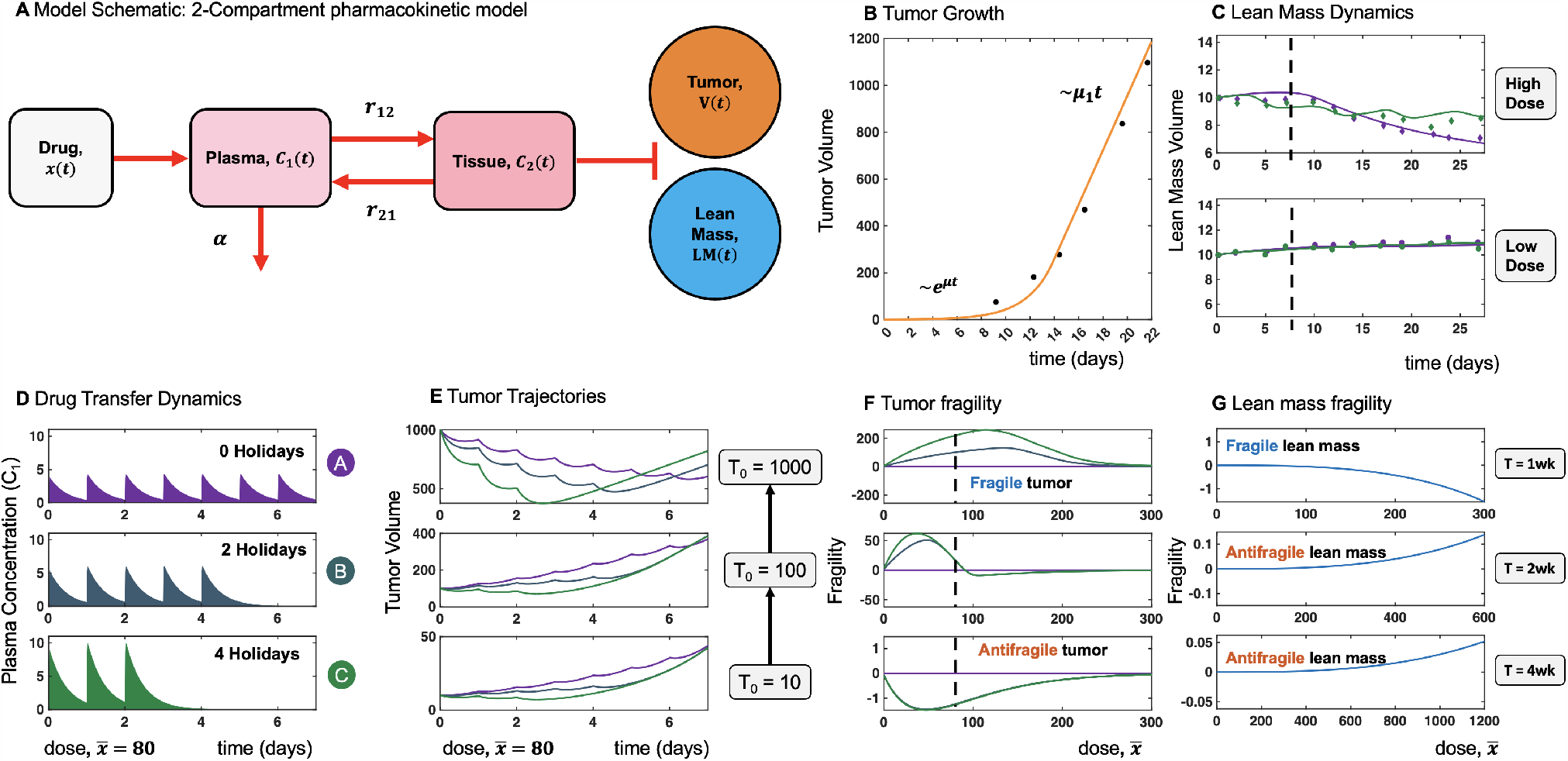
Antifragility in a 2-Compartment model with pharmacokinetics. (A) A schematic for the 2-compartment model, where drug is intravenously delivered to the blood plasma where decay occurs and is sent to the tissue compartment. Once in the tissue compartment, the drug affects the growth of the tumor and lean mass compartments. (B) Shows the untreated dynamics for the tumor compartment of the system, undergoing exponential-linear growth. (C) Shows experimental data and model predictions for lean mass volume and treatment schedules from Farhang-Sardroodi, et al^14^. The two purple schedules indicate daily doses of 35mg and 17mg over 28 days, while the two green schedules indicate doses of 50mg and 24mg on a 5 day-on, 2 day-off schedule. Model parameterization completed by minimizing the sum of squared errors. (D) A schematic for even and uneven bolus treatment injections across one week treatment and a cumulative drug total of 80mg. (E) Tumor trajectories with treatment schedules in (D) and initial tumor sizes of 10, 100, and 1000 mm^3^. As tumor size increases, even treatment becomes optimal. (F,G) Fragility of the tumor and lean mass compartments across a range of cumulative dose values and for repeated treatment schedules of 1, 2, and 4 weeks. For high tumor sizes, the tumor responds better to even treatment and is fragile, while uneven treatment is optimal for eliminating smaller tumors. For high tumor sizes optimal tumor elimination corresponds with lean mass preservation (even treatment). Simulations demonstrated in (D) - (G), used the global parameterization reported by Farhang-Sardroodi, et al^14^.

Data of the lean mass, repurposed from ref. 14, are shown in figure 4C with the model fitted to uneven (green) and even dosing schedules (purple). Lean mass is measured over time in response to an uneven schedule (5 days on treatment, 2 days on holiday, shown in green) and an even schedule (7 days on treatment, shown in purple). The purpose of this previous work was to design chemotherapy schedules that preserve lean mass across high dose (figure 4C, top) and low dose (figure 4C, bottom) schedules. However, both the cumulative dose delivered and the dosing variance (the number of treatment holidays) were altered in tandem, obfuscating first- and second-order effects. In contrast, we consider second-order effects in isolation and optimize the number of treatment holidays across identical cumulative dose by quantifying fragility in the model.

For example, panel D in figure 4 shows a range of even (purple) to uneven (green) schedules in the plasma compartment over time. We now examine the effect on tumor response (E, F) and lean mass (G) separately. Panel E shows these schedules’ effect on tumor volume as a function of the initial tumor sizes. For a mean dose of 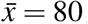, the even (no holiday) treatment schedule maximally reduces the tumor when the initial size, *T*_0_, is large.

Large tumors (*T*_0_ = 1000) tend to be very fragile (*F >* 0), while small tumors (*T*_0_ = 10) are slightly antifragile (*F <* 0). Panel F shows simulation results for tumor fragility across a range of mean dose values, with the dashed line corresponding to Panel E. Larger tumors are reduced more under schedules without holidays, while smaller tumors respond to treatments with holidays. As seen in panel F, large tumors are fragile (*F >* 0) across a large range of mean dose values, indicating the effectiveness of even treatment at reducing tumor burden. This suggests that large tumors should initially be treated with an even schedule and switched to an uneven treatment schedule at a later point in time when the tumor is sufficiently small so as to be fragile. The superiority of switching based on tumor size is shown in figure S1, illustrating the promise of fragility-guided treatment scheduling.

This model assumes no feedback of tumor size on lean mass, and thus lean mass treatment dynamics are independent of tumor size. However, lean mass has a strong dependence on time. Lean mass reduction is a function of the average exposure of the tissue to drug concentration over the last *τ* days, which ref. 14 has estimated as *τ* = 8 days (see section S4 for more details on lean mass *τ*-dependence). This time-exposure effect on lean mass leads to a shift in lean mass fragility after the first week. As seen in panel G, lean mass is initially fragile (1 week), but shifts to antifragile after 2 and 4 weeks. In the long term (>1 week), preservation of lean mass is best achieved using treatment schedules with dose holidays.

## 4 Discussion

Dose response curvature (or synonymously, antifragile theory) has been previously proposed as a graphical approach to predict outcomes based on dose response convexity and concavity^3,8^. In previous literature, this approach ignored pharmacokinetic considerations. We have constructed an extension of previous frameworks to include pharmacokinetic clearance of drug (a 1-compartment PK model). This framework allowed us to find analytical expressions for certain special cases including fast pharmacokinetic decay (*α* → ∞), no decay (*α* = 0), as well as quantifying the effectiveness of dose ordering (high or low dose first).

In general, low doses are associated with antifragility (concave dose response where uneven treatment is optimal) while high doses are associated with fragility (convex dose response where even treatment is optimal). Pharmacokinetic clearance expands the region of doses that are antifragile, but minimizes the total magnitude of antifragility.

These theoretical results build intuition to help explain chemotherapy response in a mouse-parameterized 2-compartment model. Interestingly, the dominant factor in determining fragility was the tumor size, in addition to the dose value. Large tumors are typically fragile (even is optimal) while small tumors are marginally antifragile. This implies an optimal switching point: large tumors should be treated with even therapy, until such a tumor size that is antifragile, at which point the treatment should be switched to uneven.

The modeling here also investigates the effectiveness of treatment holidays in preserving lean mass. On short time scales, the high doses of uneven treatment results in a decline in lean mass (compared to even treatment), but over longer time scales, treatment holidays prove to be a viable strategy for “resetting the clock” of the time exposure detrimental effect of chemotherapy concentration on lean mass.

Second-order effects may be driving experimental results but remain hidden due to different mean dose values, making it difficult to quantify their impact. Here we have illustrated the importance of comparing protocols with identical first-order metrics, since only then can second-order effects be properly understood. Our historic focus on mean dose and its response in cancer treatment has meant that variance in drug dose has been severely understudied. However, with the importance of dose modulation (including treatment holidays) as a means of regulating cachexia, perhaps the time for second-order effects has come.

