## Supplementary Information for "Second-order effects of chemotherapy pharmacodynamics and pharmacokinetics on tumor regression and cachexia"

### Supplemental Material

#### S1 Jensen's Inequality and Fragility

##### S1.1 Jensen's inequality

The fragility metric is motivated by Jensen's inequality, which states that for a convex function $f(x)$,

$$pf(x_1)+(1-p)f(x_2) \geq f(px_1+(1-p)x_2), \quad \forall x_1, x_2 \text{ and } p \in [0,1]. \tag{29}$$

Geometrically, this equation means that the secant line formed by the points $f(x_1)$ and $f(x_2)$ will be higher than any output $f(x)$ generated between the points $x_1$ and $x_2$. To represent cancer treatment, $f(x)$ may denote the tumor volume following a drug dose of $x$ concentration. In the case of an arbitrary number of treatment days, the generalized form of Jensen's inequality is used:

$$\sum_{i=1}^{n} p_i f(x_i) \geq f\left(\sum_{i=1}^{n} p_i x_i\right), \quad 0 \leq p_i \leq 1, \sum_{i=0}^{n} p_i = 1. \tag{30}$$

##### S1.2 Naive Fragility

Suppose the growth of a tumor only dependent on the dose $x$ of treatment and is fit by the exponential model

$$V(x) = V_0 e^{\gamma(x)}. \tag{31}$$

Here, $V_0$ denotes the initial tumor volume and $\gamma(x)$ is the drug-induced-proliferation. Now consider delivering $N$ different doses in symmetric time intervals with dosing concentrations $x_i$. The final tumor volume $V_1$ following this treatment schedule will be

$$V_\text{U} = V_0 e^{\frac{1}{N}[\gamma(x_1)+\gamma(x_2)+\ldots+\gamma(x_N)]}. \tag{32}$$

The tumor volume $V_1$ generally includes dosing schedules $\{x_i\}$ which are not identical. If these doses are delivered with the same concentration $\bar{x}$, then the final tumor volume $V_2$ will instead be

$$V_\text{E} = V_0 e^{\gamma(\bar{x})}, \quad \bar{x} = \sum_{i=1}^{N} \frac{x_i}{N}. \tag{33}$$

We call $V_\text{U}$ the tumor volume resulting from *uneven* treatment and $V_\text{E}$ the tumor volume resulting from *even* treatment. Even treatment is optimal if

$$V_0 - V_\text{U} < V_0 - V_\text{E}, \tag{34}$$

$$\Rightarrow \sum_{i=1}^{N} \frac{1}{N}\gamma(x_i) > \gamma\left(\sum_{i=1}^{N} \frac{x_i}{N}\right). \tag{35}$$

This condition mirrors Jensen's inequality in equation 30 with $f = \gamma$ and $p_i = \frac{1}{N}$. Equivalently, we expect

$$\sum_{i=1}^{N} \gamma(x_i) > N\gamma(\bar{x}) \tag{36}$$

if even treatment is optimal. This shows that if $\gamma$ is a convex dose response curve, then even treatment will be optimal. We utilize this fact to create a metric whose sign indicates whether even or uneven treatment is prefereble. The metric, fragility, characterizes this difference in performance and is defined as

$$F(\vec{X}) = \sum_{i=1}^{N} \gamma(x_i) - N\gamma(\bar{x}), \quad \vec{X} = [x_1, x_2, ..., x_N], \tag{37}$$

where $\vec{X}$ simply denotes the dosing schedule considered. It's sign will be be positive if $\gamma(x)$ is convex (even optimal) and negative if $\gamma(x)$ is concave (uneven optimal). Note that this equation does not take into account residual effects, i.e. leftover drug concentration remaining from previous dosing events. Therefore, the the metric is expected to be unreliable in cases where residual dose concentration is not negligible.

### S2 Model derivation (pharmacodynamics only)

We now consider the case when tumor volume is time dependent and determined by the ODE

$$\dot{V} = V\gamma(x(t)). \tag{38}$$

This equation may be solved analytically by inserting an appropriate $x(t)$ into the tumor volume equation. Here we wish to determine the final tumor volume following a treatment cycle of period $T$ described as

$$x(t) = \begin{cases} \bar{x} + \sigma, & 0 < t < \frac{T}{2}, \\ \bar{x} - \sigma, & \frac{T}{2} < t < T. \end{cases} \tag{39}$$

Substitution and integration can proceed to give

$$\int_{V_0}^{V} \frac{dV'}{V'} = \int_0^{\frac{T}{2}} \gamma(x+\sigma)dt + \int_{\frac{T}{2}}^{T} \gamma(x-\sigma)dt, \tag{40}$$

$$\Rightarrow \ln\left(\frac{V}{V_0}\right) = \frac{T}{2}\left((\gamma(x+\sigma) + \gamma(x-\sigma)\right), \tag{41}$$

$$\Rightarrow V(x,\sigma) = V_0 \exp\left(\frac{T}{2}\left((\gamma(x+\sigma) + \gamma(x-\sigma)\right)\right). \tag{42}$$

With this expression for the final tumor volume after a treatment cycle, the fragility $F$ can be defined as the difference in tumor volume between uneven and even treatments after a time interval

$T$. We set

$$F(x,\sigma) = V(x,\sigma) - V(x,\sigma=0), \tag{43}$$

$$\Rightarrow F(x,\sigma) = V_0 \left[ \exp\left( \frac{T}{2} \left( (\gamma(x+\sigma) + \gamma(x-\sigma) \right) \right) - \exp\left(T\gamma(x)\right) \right]. \tag{44}$$

Finding the antifragile-fragile boundary generally requires solving for $x_B$ such that $F(x_B) = 0$. Substitution and simplification gives

$$\frac{1}{2}\left(\gamma(x_B+\sigma) + \gamma(x_B-\sigma)\right) - \gamma(x_B) = 0. \tag{45}$$

Note that when the left hand side is greater than or equal to zero, Jensen's inequality is recovered from Equation 29 with $f(x) = \gamma(x)$, and $x_i = \bar{x} \pm \sigma$.

The dose response $\gamma(x)$ can be expressed as the Hill function $H(x)$, an empirical pharmacodynamic equation, with an appropriate choices of parameters. It is given by

$$H(x) = E_0 + \frac{x^n(E_1 - E_0)}{x^n + C^n}. \tag{46}$$

Setting $\gamma(x) = H(x)$, the above equation becomes

$$\frac{1}{2}\left(H(x_B+\sigma) + H(x_B-\sigma)\right) - H(x_B) = 0, \tag{47}$$

which is precisely the expression determined by West, et al[8]. Though this antifragile-fragile boundary generally has a small dependence on $\sigma$, the paper also showed that for small $\sigma$,

$$x_B = \left(\frac{n-1}{n+1}\right)^{1/n}. \tag{48}$$

So in the case of small $\sigma$, the antifragile-fragile boundary is only dependent on parameters present in the drug-induced proliferation.

### S3 1-Compartment Model Analysis

#### S3.1 Analytical solution to 1-compartment model

The complete 1-compartment model used is given by:

$$\dot{V} = V\gamma(x(t)), \tag{49}$$

$$\gamma(x) = H(x) = E_0 + \frac{x^n(E_1 - E_0)}{x^n + C^n}, \tag{50}$$

$$\dot{x} = -\alpha x + \sum_i d_i \delta(t - t_i), \tag{51}$$

where $\delta$ is the Dirac-delta function. To continue, we attempt to solve for the dose $x(t)$ independently with Laplace transforms:

$$\mathrm{L}[\dot{x}] = \mathrm{L}\left[-\alpha x + \sum_i d_i \delta(t - t_i)\right], \tag{52}$$

$$\mathrm{L}[\dot{x}] = -\alpha \mathrm{L}[x] + \sum_i d_i \mathrm{L}[\delta(t - t_i)], \tag{53}$$

where the second line follows because the Laplace transform is linear. Using known properties of the Laplace transform, equation 53 becomes

$$s\mathrm{L}[x] - x(0) = -\alpha \mathrm{L}[x] + \sum_i d_i e^{-t_i s}. \tag{54}$$

Solving for $\mathrm{L}[x]$ gives

$$\mathrm{L}[x] = \frac{x(0)}{s+\alpha} + \frac{1}{s+\alpha}\sum_i d_i e^{-t_i s}. \tag{55}$$

For convenience we document the following useful identities for the Laplace transform:

$$\mathrm{L}\left[e^{-ct}\right] = \frac{1}{s+c}, \tag{56}$$

$$\mathrm{L}\left[e^{-c(t+t_0)}\right] = \mathrm{L}\left[e^{-ct}e^{-ct_0}\right] = \frac{1}{s+c}e^{-ct_0}, \tag{57}$$

$$\mathrm{L}[u_c(t)g(t)] = e^{-cs}\mathrm{L}[g(t+c)], \tag{58}$$

where $u_c(t)$ is the step function

$$u_c(t) = \begin{cases} 0 & t < c, \\ 1 & c \leq t. \end{cases} \tag{59}$$

These identities alongside equation 55 provide an expression for the drug dose as a function of time:

$$x(t) = x(0)e^{-\alpha t} + \sum_i d_i u_{t_i}(t) e^{-\alpha(t-t_i)}. \tag{60}$$

In the case of two doses only, we let

$$x(0) = x_0 + \sigma, \tag{61}$$

$$d_1 = x_0 - \sigma, \tag{62}$$

$$t_1 = \frac{T}{2} \tag{63}$$

$$d_j = 0, \quad j > 1 \tag{64}$$

which implies

$$x(t) = (x_0 + \sigma)e^{-\alpha t} + (x_0 - \sigma)u_{t_1}(t)e^{-\alpha(t-\frac{T}{2})}. \tag{65}$$

Note that if $(x_0+\sigma)e^{\alpha\frac{T}{2}} \approx 0$, then

$$x(t) = \begin{cases} (x_0+\sigma)e^{-\alpha t}, & 0 < t < \frac{T}{2}, \\ (x_0-\sigma)e^{-\alpha\left(t-\frac{T}{2}\right)}, & \frac{T}{2} < t < T, \end{cases} \tag{66}$$

which is the fast-PK approximation. To find the final tumor volume under this approximation at time $T$, we substitute equation 66 into 49 and integrate:

$$\int_{V_0}^{V} \frac{dV'}{V'} = \int_0^{\frac{T}{2}} H\left((x+\sigma)e^{-\alpha t}\right) dt + \int_{\frac{T}{2}}^{T} H\left((x-\sigma)e^{-\alpha\left(t-\frac{T}{2}\right)}\right) dt. \tag{67}$$

Both integrals have the form

$$I = \int \left[E_0 + \frac{x^n e^{-\alpha n t}(E_1-E_0)}{x^n e^{-\alpha n t} + C^n}\right] dt, \tag{68}$$

and this can be solved by performing a $u$-substitution with $u = x^n e^{-\alpha n t} + C^n$. The integral becomes

$$I = E_0 t - \frac{E_1-E_0}{\alpha n} \int \frac{1}{u} du, \tag{69}$$

$$\Rightarrow I = E_0 t - \frac{E_1-E_0}{\alpha n} \ln\left(x^n e^{-\alpha n t} + C^n\right) \tag{70}$$

up to a constant. Altogether, the formula for the tumor volume at time $t = T$ is

$$V(t=T) = V_0 \exp\left(E_0 T - \frac{E_1-E_0}{\alpha n}\left[\ln\left(\frac{(x+\sigma)^n e^{-\alpha n \frac{T}{2}} + C^n}{(x+\sigma)^n + C^n}\right) + \ln\left(\frac{(x-\sigma)^n e^{-\alpha n \frac{T}{2}} + C^n}{(x-\sigma)^n + C^n}\right)\right]\right) \tag{71}$$

or

$$V(t=T) = V_0 \exp K(\sigma, \alpha, T) \tag{72}$$

as expressed in the paper. Noting the definition of Fragility, we can see that the antifragile-fragile boundary can be determined by the implicit formula:

$$\ln\left(\frac{(x_B+\sigma)^n e^{-\alpha n \frac{T}{2}} + C^n}{(x_B+\sigma)^n + C^n}\right) + \ln\left(\frac{(x_B-\sigma)^n e^{-\alpha n \frac{T}{2}} + C^n}{(x_B-\sigma)^n + C^n}\right) - 2\ln\left(\frac{x_B^n e^{-\alpha n \frac{T}{2}} + C^n}{x_B^n + C^n}\right) = 0, \tag{73}$$

which has no analytical solution.

#### S3.2 Limit 1: $\alpha \to \infty$

We seek the antifragile-fragile boundary of the 1-compartment model under limiting cases. The general problem involves finding $x_B$ such that:

$$F(x, \sigma, \alpha) = V_0\left[\exp K(\sigma, \alpha, T) - \exp K(0, \alpha, T)\right] = 0. \tag{74}$$

Using equation 73 and first considering the fast-PK limit $\alpha \to \infty$, $x_B$ is seen to satisfy

$$\ln\left(\frac{C^{2n}}{\left[C^n + (x_B+\sigma)^n\right]\left[C^n + (x_B-\sigma)^n\right]}\right) = \ln\left(\frac{C^{2n}}{C^{2n} + 2C^n x_B^n + x_B^{2n}}\right). \tag{75}$$

Rearrangement gives our final implicit solution:

$$x_B^{2n} + 2C^n x_B^n - C^n\left[(x_B-\sigma)^n + (x_B+\sigma)^n\right] - (x_B-\sigma)^n(x_B+\sigma)^n = 0. \tag{76}$$

#### S3.3 Limit 2: $\alpha \to 0$

The second limiting case involves finding $x_B$ when $\alpha \to 0$. This can be found by explicitly taking a limit, but this requires L' Hopital's rule and several lines of algebra due to the $\alpha^{-1}$ term in equation 71. Instead, setting $\alpha = 0$ in equation 66 and integrating, we get

$$\int_{V_0}^{V} \frac{dV'}{V'} = \int_0^{\frac{T}{2}} H(x+\sigma)dt + \int_{\frac{T}{2}}^{T} H(x-\sigma)dt, \tag{77}$$

$$\Rightarrow \ln\left(\frac{V}{V_0}\right) = \frac{T}{2}\left((H(x+\sigma)+H(x-\sigma)\right), \tag{78}$$

$$\Rightarrow V(t=T) = V_0 \exp\left(\frac{T}{2}\left((H(x+\sigma)+H(x-\sigma)\right)\right). \tag{79}$$

The corresponding definition of Fragility is

$$F(x,\sigma) = V(x,\sigma) - V(x,\sigma=0), \tag{80}$$

$$\Rightarrow F(x,\sigma) = V_0\left[\exp\left(\frac{T}{2}\left((H(x+\sigma)+H(x-\sigma)\right)\right) - \exp\left(TH(x)\right)\right]. \tag{81}$$

This implies

$$\frac{1}{2}\left(H(x_B+\sigma)+H(x_B-\sigma)\right) - H(x_B) = 0, \tag{82}$$

which reproduces the result present in section S2. Therefore, when $\alpha \to 0$, the phamacokinetic model becomes the pharmacodynamic-only model.

#### S3.4 Residual concentration

If the drug concentration from previous treatment is not negligible when another dose is delivered, then the order of delivery affects the average drug concentration in the tumor. We show this by considering a dosing schedule similar to equation 65, now with

$$x(t) = Ae^{-\alpha t} + Bu_{t_1}(t)e^{-\alpha(t-\frac{T}{2})}. \tag{83}$$

This represents the pharmacokinetics of a treatment schedule with drug delivery concentrations $\{A,\ B\}$. The time-averaged drug concentration $\langle x\rangle$ can be determined with

$$\langle x\rangle = \frac{1}{T}\int_0^T x(t)dt. \tag{84}$$

By substituting equation 83 into equation 84, we find that

$$\langle x\rangle = \frac{1}{T}\left(\int_0^T Ae^{-\alpha t}dt + \int_0^T Bu_{\frac{T}{2}}(t)e^{-\alpha(t-\frac{T}{2})}dt\right), \tag{85}$$

$$\Rightarrow \langle x\rangle = \frac{1}{T\alpha}\left(A+B-\left(Ae^{-\alpha\frac{T}{2}}+B\right)e^{-\alpha\frac{T}{2}}\right). \tag{86}$$

Subtracting the average concentration resulting from schedules $\{A,\ B\}$ and $\{B,\ A\}$ demonstrates a non-zero difference in drug concentrations:

$$\Delta = \frac{e^{-\alpha\frac{T}{2}}}{T\alpha}(A-B)\left(1-e^{-\alpha\frac{T}{2}}\right). \tag{87}$$

If $A > B$, then $\Delta > 0$, which implies the schedule $\{A,\ B\}$ delivers more drug in a time interval $T$. This would correspondingly eliminate more tumor and imply that delivering a higher first dose is preferable for the patient. For $A = \bar{x}+\sigma$ and $B = \bar{x}-\sigma$, the difference in average drug concentrations between schedules is

$$\Delta = \frac{e^{-\alpha\frac{T}{2}}}{T\alpha}2\sigma\left(1-e^{-\alpha\frac{T}{2}}\right). \tag{88}$$

### S4 The 2-Compartment Cachexia Model

The entire model by Farhang et al[14] is:

$$\frac{dC_1}{dt} = k_{21}C_2(t)\frac{V_2}{V_1} - k_{12}C_1(t) - k_{10}C_1(t) + \frac{1}{V_1}\sum_i d_i\delta(t-t_i) \tag{89}$$

$$\frac{dC_2}{dt} = k_{12}C_1(t)\frac{V_1}{V_2} - k_{21}C_2(t) \tag{90}$$

$$\frac{dT}{dt} = \mu_0 T\left(1+\left(\frac{\mu_0}{\mu_1}T\right)^{\eta}\right)^{-\frac{1}{\eta}} - \kappa C_2 T \tag{91}$$

$$p(t) = p_0 + \frac{p_1}{1+\frac{M(t)}{m}} \tag{92}$$

$$v(t) = v_0 + \frac{v_1}{1+\frac{M(t)}{m}} \tag{93}$$

$$\overline{C}_2(t) = \frac{1}{\tau}\int_{t-\tau}^{t} C_2(s)ds \tag{94}$$

$$\frac{dS}{dt} = \left(1-\left(\frac{\overline{C}_2(t)}{R_d}\right)^3\right)(2p(t)-1)v(t)S \tag{95}$$

$$\frac{dM}{dt} = \left(1-\left(\frac{\overline{C}_2(t)}{R_d}\right)^3\right)2(1-p(t))v(t)S - d_0M. \tag{96}$$

The concentrations $C_1$ and $C_2$ represent the drug present in the plasma and tissue, respectively. These inhibit the growth dynamics of a tumor $T$ while also eliminating stem ($S$) and muscle ($M$) cells composing lean mass ($L = S+M$). The parameter set provided by 14 is used unless otherwise mentioned.

#### S4.1 Fragility guides treatment scheduling

As seen in the main text (figure 4E), large tumors are typically fragile (even is optimal) while small tumors are slightly antifragile (uneven is optimal). This leads to the hypothesis that large tumors should initially be treated with an even schedule and switched to an uneven treatment schedule at a later point in time when the tumor is sufficiently small so as to be fragile.

An example is shown in figure S1, comparing two weeks of even (purple) and uneven (green) treatment to a well-timed switching strategy of one week even followed by one week uneven (light blue). Due to an initial large, fragile tumor the even treatment strategy is initially outperforms the uneven strategy (note the differences at the halfway point marked with a dashed line). Subsequently, as the tumor size approaches antifragility, then a switch to uneven therapy (blue line) is optimal. We note that gains by switching are modest, due to the small values of antifragility calculated in figure 4F (bottom panel).

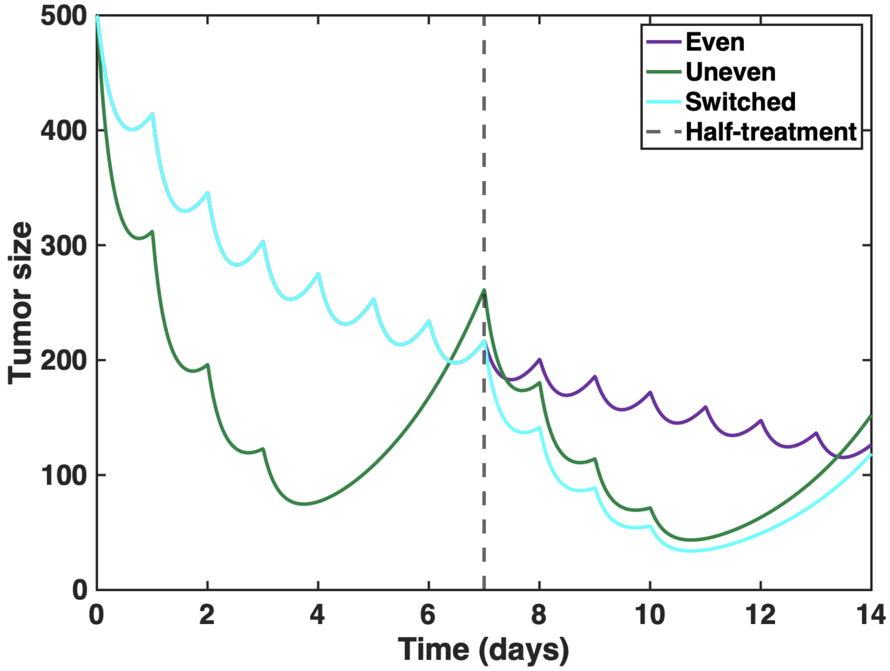


**Figure S1. Fragility guides treatment scheduling** Using the model shown in figure 4, we compare treatment schedules over two weeks: even (purple), uneven (green) and switching (blue). The switching strategy (1 week even followed by 1 week uneven) outperforms even and uneven treatment. Parameterization: simulation is performed using the global parameterization reported by Farhang-Sardroodi, et al[14].
